# The polymorphic PolyQ tail protein of the Mediator Complex, Med15, regulates variable response to stress

**DOI:** 10.1101/652669

**Authors:** Jennifer E.G. Gallagher, Casey Nassif, Amaury Pupo

## Abstract

The Mediator is a multi-protein complex composed of subunits called head, body, tail, and CDK that is conserved from yeast to humans and plays a central role in transcription. However, not all the components are required for basal transcription. Components of the tail are not essential but to varying degrees are required for growth in different stresses. While some stresses are familiar such as heat, desiccation, and starvation, others are exotic, yet yeast can elicit a successful stress response. MCHM is a hydrotrope that induces growth arrest in yeast. By exploiting genetic variation, specifically in Med15, between yeast strains, we found that a naturally occurring Med15 allele with polyQ (polyglutamine) expansion conferred MCHM sensitivity. Expansion in polyQ repeat can induce protein aggregation and in humans can cause neurodegenerative diseases. In yeast, the MCHM sensitivity was not from a loss of function as the reciprocal hemizygous hybrids were all sensitive and the homozygous null mutant was less sensitive than the hemizygous hybrids. This suggests that there is an incompatibility between Mediator components from genetic divergent yeast strains. Transcriptomics from yeast expressing the incompatible Med15 (longer polyQ repeats in the strain with fewer repeats) changed gene expression in diverse pathways. Med15 protein existed in multiple isoforms, mostly from likely post-translational modifications and different alleles have different patterns of isoforms. Stability of both alleles of Med15 was dependent on Ydj1, a J-type chaperone. The protein level of the incompatible Med15 allele was lower than the compatible allele and was turned over faster. Med15 is tethered to the rest of the Mediator complex via Med2 and 3. Deletion of either Med2 or Med3 changed the Med15 isoform patterns in a similar manner. Whereas deletion of Med5, a distal component of the Mediator tail, did not change the pattern. The *med2* and *med3* mutants were similarly sensitive to MCHM while *med5* mutants were not. Differences in the phenotype of yeast carrying different Med15 alleles extend to other stresses. The incompatible allele of Med15 improved growth of yeast to chemicals that produce free radicals and the compatible allele of Med15 improved growth to reducing agents, caffeine, and hydroxyurea. Med15 directly interacts with Gcn4 and other TFs and *in vitro* form phase-separated droplets. This variation may reflect the positive and negative role that Med15 has in transcription. Genetic variation in transcriptional regulators can magnify differences in response to environmental changes, in contrast, genetic variation in a metabolic enzyme. These polymorphic control genes are master variators.

## Background

Changing the transcriptional landscape is a key step in reorganizing cellular processes in response to stress. RNA polymerase II (pol II) transcription is regulated in a stress-specific manner by multiple post-translational modifications and a host of transcription factors (TFs). These transcription factors do not interact directly with pol II and general transcription factors (GTFs), together called the pre-initiation complex, but through a multi-protein complex called the Mediator. The Mediator itself is composed of four domains: the head, body, tail and kinase domains (Figure 1A). The head interacts with pol II and GTFs while the tail interacts with specific TFs (reviewed (1)). The tail is composed of Med2, Med3 (Pgd1), Med5 (Nut1), Med15 (Gal11) and Med16 (Sin4) and the C-terminal end of Med14 connects the tail with the body of the Mediator complex (Figure 1B, (Tsai *et al.* 2014, 2017; Robinson *et al.* 2015)). The tail is the most diverged between species and binding of TF changes the confirmation (5).

**Figure 1.**
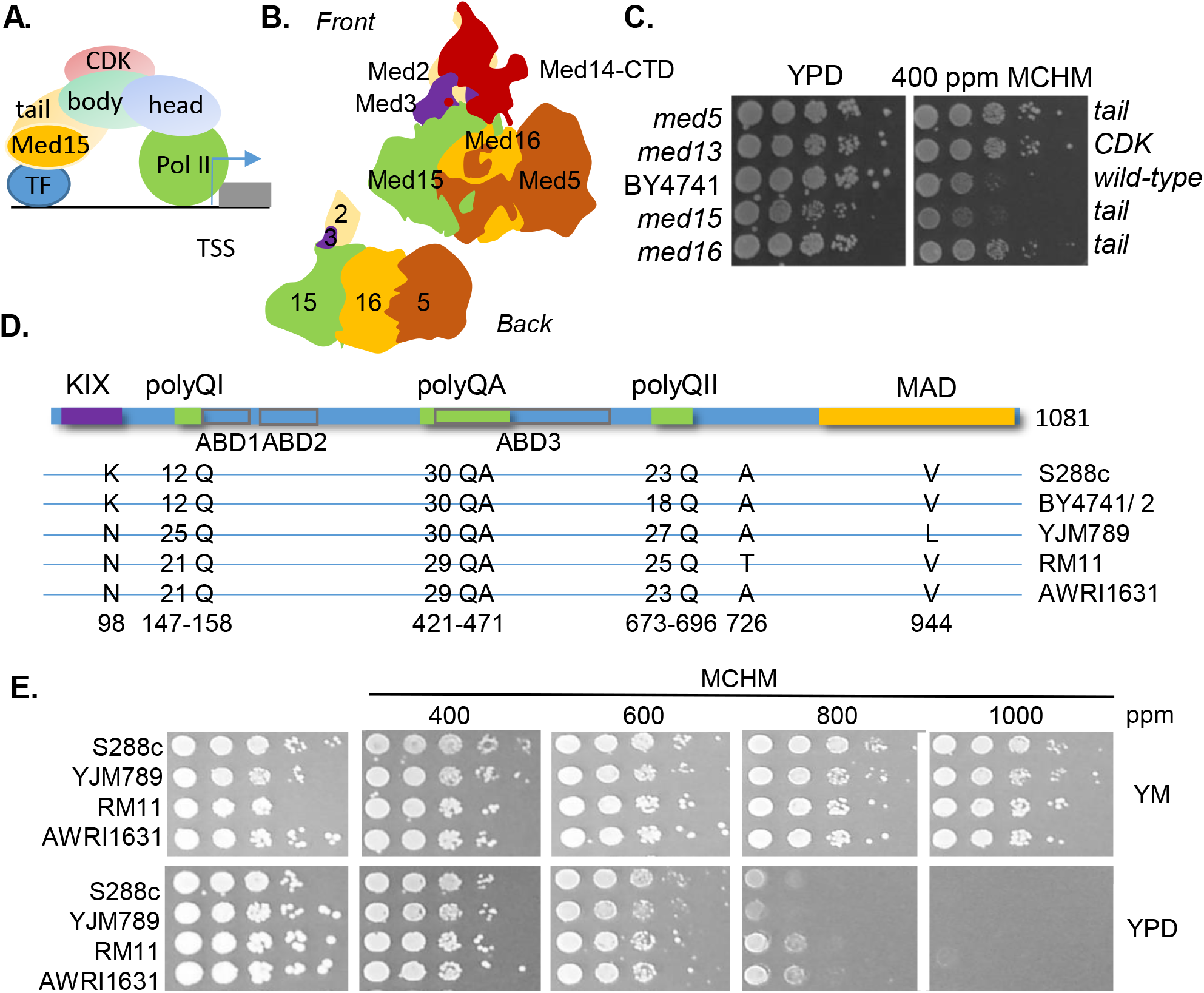
Role of Mediator tail in response to MCHM **A.** Schematic of the Mediator complex. Med15 as part of the tail subcomplex directly interacts with transcription factors (TF). The body of the Mediator complex tethers the CDK (cyclin-dependent kinase). The head directly interacts with RNA polymerase II (Pol II) at promoter regions to initiation transcription at transcriptional start sites (TSS) of genes (gray box). **B.** Representation of protein components of the Mediator tail based on structures and modeling (Robinson *et al.* 2015). Med2 (beige), Med14-CTD (red), Med3 (purple), Med15 (green), Med16 (orange) and Med5 (brown) comprise the tail of the Mediator complex. From the back view, Med5, Med16, Med15, Med3 and Med2 (in order from farthest to closest to the rest of the Mediator complex) associate with Med14 (not pictured here). **C.** Growth assays of yeast with different components of the Mediator tail knocked out in BY4741 grown with and without 400 ppm MCHM in YPD. **D.** Schematic of Med15 protein. Above the blue line are polymorphic domains including the KIX domain, polyglutamine domain I (polyQI), polyglutamine/ alanine domain (polyQA), polyglutamine domain II (polyQII) and the Mediator activation/ association domain (MAD). Under the blue line are the fuzzy domains represented as ABD1-3 (activator-binding domains) in gray outlined boxes (Herbig *et al.* 2010; Brzovic *et al.* 2011). The Med15 polymorphic amino acids are diagramed below from five genetically diverse yeast. Amino acid numbers are based on S288c. **E.** Growth assays of genetically diverse yeast strains in the presence of MCHM on different growth media with increasing concentrations of MCHM. Yeast were spotted in ten-fold dilutions onto minimal media supplemented with lysine (YM), or rich media (YPD). RM11, S288c (GSY147), AWRI1631 are MATa prototrophs while YJM789 is a MATalpha *lys2* strain.

Med15 has a curious amino acid sequence (reviewed (6)). Med15 from a common lab strain S288c is 16% glutamines and 11% asparagines. Two regions have two polyglutamine (polyQ) tracts separated by a polyQA. These regions along with the KIX domain interact with TF through multiple sites termed fuzzy domains which are intrinsically disordered regions. Med15 makes multiple contacts with Gcn4 including polymorphic polyQ repeats and KIX domain (Jedidi *et al.* 2010). Overexpression of Med15 causes protein aggregation, presumably via the polyQ and polyQA regions of this region alone aggregates in response to hydrogen peroxide (Zhu *et al.* 2015). Overexpression of the first polyQ and polyQA of Med15 reduces cell growth in unstressed cells and salt exposed yeast but rescues growth in the presence of rapamycin (8). Full-length Med15 also forms cytosolic foci in yeast exposed to hydrogen peroxide (8). The pathogenic effects of polyQ proteins were uncovered when the causative mutation for Huntington’s disease was discovered (9). Huntington’s disease causes progressive neurodegeneration in people who inherit a single copy of HTT with the polyQ expansion inducing protein aggregation (reviewed (10)). Aggregation of polyQ expansion proteins in yeast can be reduced by overexpression of heat shock chaperone proteins (11).

Ydj1 is a highly expressed general type I Hsp40 protein (J-type) chaperone that localizes to the mitochondria, cytoplasm, and nucleus. Yeast lacking Ydj1 function are sensitive to multiple classes of chemicals (Gillies *et al.* 2012). Hsp40 proteins work with Hsp70 to refold misfolded protein or target them for degradation. They also have roles in translation, translocation across membranes and conformation changes induced by amyloid fibrils. Overexpression of Ydj1 can cure prions (Hines *et al.* 2011). Prions are a group of proteins that not only aggregate but also can induce the aggregation of natively folded proteins. Prions can cause contagious neurodegenerative diseases in humans and switches in the prion state to provide epigenetic plasticity in phenotypic response to stresses by regulating the enzymatic function (14). When overexpressed Med15 will aggregate *in vivo* (8). Overlapping the polyQ domains are the intrinsically disordered regions (IDR) that form fuzzy interactions with TFs, in particular, Gcn4 (15–18). An N-terminal fragment of Med15 containing the first polyQ and the polyQA domain will form liquid phase condensates also known as liquid-liquid phase separation with Gcn4 at low *in vitro* (19). These condensates are dynamic and behave like a liquid (reviewed (20)). A mutant of Gcn4 which forms liquid droplets (phase separation) condensates with Med15, no longer activates transcription (19). The transition from single phase to liquid phase droplet increases the local concentration of factors by forming non-membrane found compartments that flow and fuse with surface tension (reviewed in (21)). Liquid phase droplets can be induced by chemicals and act as compound concentrators. IDR interactions may be a more general mechanism to increase the local concentration of proteins within liquid droplets, changing protein confirmations, and adding complexity regulating cellular metabolism and environmental responses.

MCHM is a coal-cleaning chemical that acts as a hydrotrope *in vitro* (22). Hydrotropes increase the solubility of organic compounds by inducing liquid phase condensates. Currently, hydrotropes are not considered detergents and detergents function at lower concentrations to solubilize compounds. ATP is a biological active hydrotrope (23–25) and RNA can induce changes in solubility of Whi3 via phase separation (26). These types of hydrotropes act to concentrate compounds. Exposure to MCHM induced growth arrest in yeast by changing a wide range of biochemical pathways including ionome (22) and amino acids (27). The Mediator binds upstream of many genes across pathways, including stress responsive genes. Numerous studies have explored the role of Med15 via knockouts on microarrays and later RNA-seq. Removing the entire coding region not only removes the function of a protein but also alters the structure of complexes containing that protein. Gene knockouts are rarely found in nature while, indels, copy number variation, and SNPs are the most common mutations. By assessing the role of naturally variable protein, the integrity of the Mediator is maintained and the specific function of Med15 can be addressed in response to hydrotropic chemicals such as MCHM.

As the altered state of protein conformation/ phase (single verse liquid) are coming to light, the highly variable Med15 was further characterized. Polymorphic proteins that regulate gene expression are likely to allow a small genetic variation to have a large impact on phenotypic variation. A single polymorphism of threonine to isoleucine removed potential phosphorylation in Yrr1, a transcription factor, confers 4NQO sensitivity but has the benefit of increased respiration (28, 29). These polymorphic proteins are termed master variators (28). MCHM is a hydrotrope that increases protein solubility (22) and Med15 can exist as liquid droplets *in vivo* (19). Genetic variation of Med15 regulated cellular response to MCHM but not at the protein level or isoform patterns. Replacing the BY4741 yeast strain Med15 with the Med15 from the parental strain S288c or a highly divergent strain, YJM789, uncovered an incompatibility between subunits. In a hybrid strain, the hemizygous strain with either allele of Med15 was more sensitive to MCHM than the homozygous diploid. Two different alleles of Med15 protein displayed multiple isoforms which likely represent posttranslational modifications. Protein level and isoform stability were dependent on a protein chaperone, Ydj1. When proteins that tether Med15 to the complex were knocked out, the pattern of the incompatible Med15^YJM789^ allele shifted to be more like the compatible Med15 allele. The incompatible Med15^YJM789^ in MCHM did not act as a null allele and in other stress conditions, such as exposure to 4NQO and hydrogen peroxide that generate free radicals, yeast carrying the Med15^YJM789^ have increased growth over the strain with Med15^S288c^. Med15 is a master variator that provides phenotypic plasticity.

## Results

The growth of yeast with different components of the Mediator complex knocked out were tested in the response to MCHM (Figure 1C). As the tail directly interacts with the TFs *med15, med16*, and *med5* knockouts were tested and *med13* from the CDK was chosen because it is on the other side of the complex from the tail. Mutants in *med5, med13*, and *med16* grew better than wild-type yeast (BY4741) in response to MCHM than the parental strain, BY4741, while the growth of the *med15* mutant was inhibited after three days of growth. Med15 is a 120 kDa protein with multiple protein domains (Figure 1D). The KIX domain is at the N-terminal domain and interacts with TFs. Two polyQ tracts are separated by a polyQA tract. Between species, C-terminal end of the Mediator is highly divergent and required for the association to the Mediator complex, Mediator activation/ association domain (MAD). MAD is heavily phosphorylated but the exact roles of these phosphorylations have not been determined (Albuquerque *et al.* 2008; Holt *et al.* 2009; Soulard *et al.* 2010; Swaney *et al.* 2013). Between polyQ I and polyQ II and partially overlapping with the polyQA tract are three ABD (Activator Binding Domains) regions (Pacheco *et al.* 2018). The ABDs and KIX domain form fuzzy interactions with TFs (Warfield *et al.* 2014; Tuttle *et al.* 2018). The unusual structure of Med15 leads us to investigate either Med15 from other strains had genetic variation in the polyQ tracts. Med15 from five genetically diverse yeast had between 12 and 25 Qs in the polyQ I and between 18 and 27 Qs in polyQII. The polyQA only differed by one less QA repeat in RM11 and AWRI1631 Med15 alleles. There were three other non-synonymous SNPs were K98N, A726T, and V944L using S288c numbering. These strains were then tested on increasing concentrations of MCHM in YPD and YM (yeast minimal media with no amino acids, Figure 1E). No decrease of growth in the strains was detected in YM at the highest concentration of 1000 ppm MCHM which is the limit of solubility of MCHM in media. These strains are more robust than BY4741 and growth was slowed at 800 ppm MCHM in YPD with YJM789 being the most sensitive.

YJM789 and BY4741 were selected for further study because their alleles of Med15 represent the variation in polyQ lengths, differences in MCHM resistance and available genetic markers. Reciprocal hemizygosity assays were carried out. *MED15* was knocked out in haploid parent strains and diploids selected. Both the *MED15*^*YJM789*^*/Δ* and the *MED15*^*BY*^*/Δ* diploids were equally sensitive to MCHM (Figure 2A). However, when compared to the homozygous mutant, the hemizygotes were more sensitive. Suggesting there is no impact of the different alleles of Med15 on MCHM response in the context of a hybrid Mediator complex, but Med15 is important possibly as a gene dosage effect in respect to the stoichiometry of the Mediator complex.

**Figure 2.**
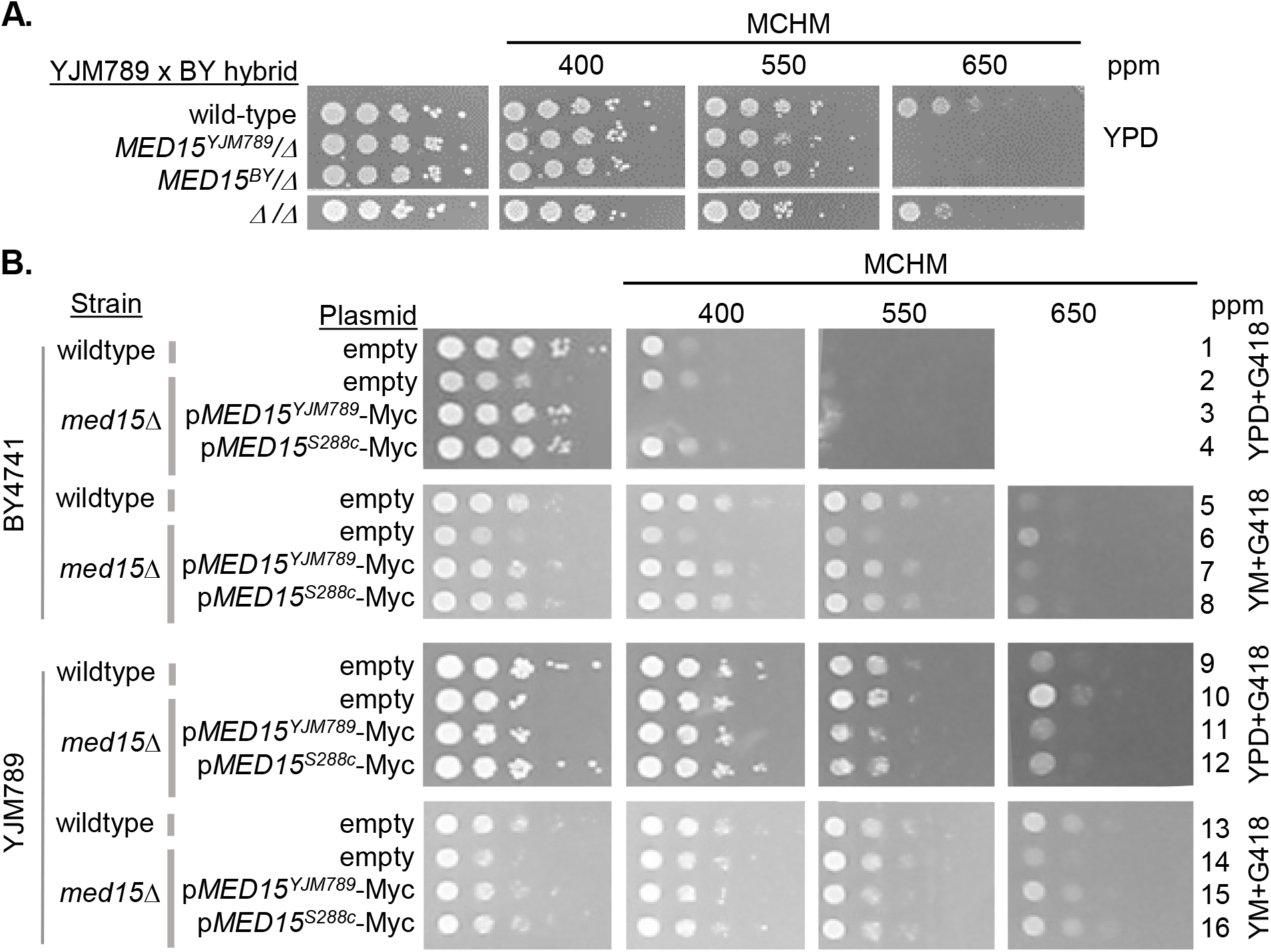
Genetic variation in Med15 contributes to variation in MCHM response. **A.** Reciprocal hemizygotes of Med15 in BY4741xYJM789 hybrids were grown on MCHM in YPD. Med15 was tagged at the chromosomal locus with 13xMyc at the C-terminal end or knockout with a dominant drug marker in haploid parents. Yeast were then mated, and diploids selected. **B.** Med15 allele swap in BY4741 and YJM789 was carried out by cloning Med15-13xMyc with *KanR* onto pRS316. Med15 plasmids were transformed into wild-type and *med15::NatR* stains in the BY4741 and YJM789 (YJM789K5a, a MATa prototroph) backgrounds. Plasmids were maintained by growth on YPD with G418. Glutamate (MSG) was used as the nitrogen source in minimal media with histidine, uracil, leucine, and methionine to supplement BY4741 so that G418 would be selective and maintain the plasmid. The empty plasmid is pGS35 (*KanR*).

The gene dosage could mask allelic differences and to control for this, *MED15* from YJM789 and S288c strains to swap alleles in BY4741 and YJM789 haploid knockouts. *MED15* alleles were cloned from yeast which had their *MED15* alleles tagged with Myc at the chromosomal location with the KanR marker. Both alleles are expressed under their endogenous promoter from a single copy plasmid. We did find that the Myc tag on Med15 increased the sensitivity of yeast MCHM when comparing the reciprocal hemizygotes to the untagged strains (Figure S1A). *MED15* was knocked out in BY4741 and YJM789 and transformed with the two alleles of Med15 with the empty plasmid as the negative control and grown in YPD or YM (with MSG as the nitrogen source instead of ammonium sulfate). Wildtype BY4741 grew slower than BY4741 carrying the S288c allele of Med15 in the YPD with low levels of MCHM for two days (Figure 2B row 1 and 4). In these same conditions, there is very little change in the growth of the BY4741 *med15* knockout (Figure 2B row 2). However, the yeast with Med15^YJM789^ was severely affected by MCHM. Consistent with the growth of the other strains, more MCHM was required in YM to slow growth of yeast and there was no difference between the three alleles of Med15 (row 5,6, and 8). The *med15* knockout grew slower in YM but appeared to be not affected by MCHM when the slow growth was also taken into account (Figure 2B row 6). YJM789 growth was not affected by the alleles of Med15 expressed in YPD or YM (Figure 2B row 9, 11, 12, 13, 15, 16). YJM789 *med15* mutant grew slower in YM, yet mutant growth was about the same in 400 ppm MCHM in YPD and YM. At 550 and 650 ppm MCHM in YPD, the knockout grew better in YPD than yeast with Med15 but not in YM.

It is surprising that BY4741 Med15^YJM789^ yeast were more sensitive to MCHM in YPD than the *med15* knockout yeast. To test if the Med15^YJM789^ was a dominant negative allele, Med15^YJM789^ was expressed in wild-type BY4741 with endogenous Med15^BY^. Expressing both Med15^YJM789^ and Med15^BY^ in yeast did not change growth in YPD with MCHM and no difference was noted when compared to yeast with Med15^S288c^ and Med15^BY^ in the By4741 *med15* strain (Figure S1B row 7 and 8). However, yeast expressing both Med15^YJM789^ and Med15^BY^ were more sensitive to MCHM in MSG (YM with glutamine as the nitrogen source to maintain the KanR plasmid in minimal media) with high levels of MCHM (Figure S1B row 7 and 8).

To determine if the protein levels of Med15 contribute to differences in MCHM sensitivity, expression levels of the cloned alleles *MED15* in the allele swapped strains were measured. Med15-Myc proteins were immunoprecipitated because the levels are too low to detect by western blot without enrichment. Yeast were grown to mid-log phase in YPD or YM with amino acids supplemented and then shifted to media containing MCHM for 30 minutes. Med15^YJM789^ levels were lower than Med15^S288c^ in all conditions tested, YPD, YM with and without MCHM. In general, the levels of both alleles were lower in YM. Med15^YJM789^ levels did also appeared to decrease in YPD with MCHM but the decreased levels did not explain the MCHM sensitivity as the *med15* knockout was not as sensitive as yeast carrying the Med15^YJM789^ allele. Similarly, yeast with Med15^YJM789^ grew similarly to yeast carrying Med15^S288c^ in YM and the levels of Med15 protein were very different in YM (Figure 3A). It is also curious to note, with the Myc tagged Med15^S288c^ is predicted to be 140 kDa with pI at 6.61 and Med15^YJM789^ is predicted to be 142 kDa with a pI at 6.48. Med15^S288c^ protein runs above the 150 kDa marker as multiple bands despite being shorter than Med15^YJM789^ which runs truer to size. In part, the differences in Med15 proteins levels can be attributed to differences in mRNA levels. Global mRNA levels were quantified by Illumina sequencing of three biological replicates (Figure 3B). *MED15*^*YJM789*^ mRNA decreased in YPD with MCHM and was equivalent in YM irrespective of MCHM. The levels of *MED15*^*S288c*^ mRNA levels also tracked with protein levels. The *MED15* promoter contains 4 SNPs that were included on the plasmid which are in relation to the start codon of S288c to YJM789: A-8T, A-209G, A-365G, and T-449C.

**Figure 3.**
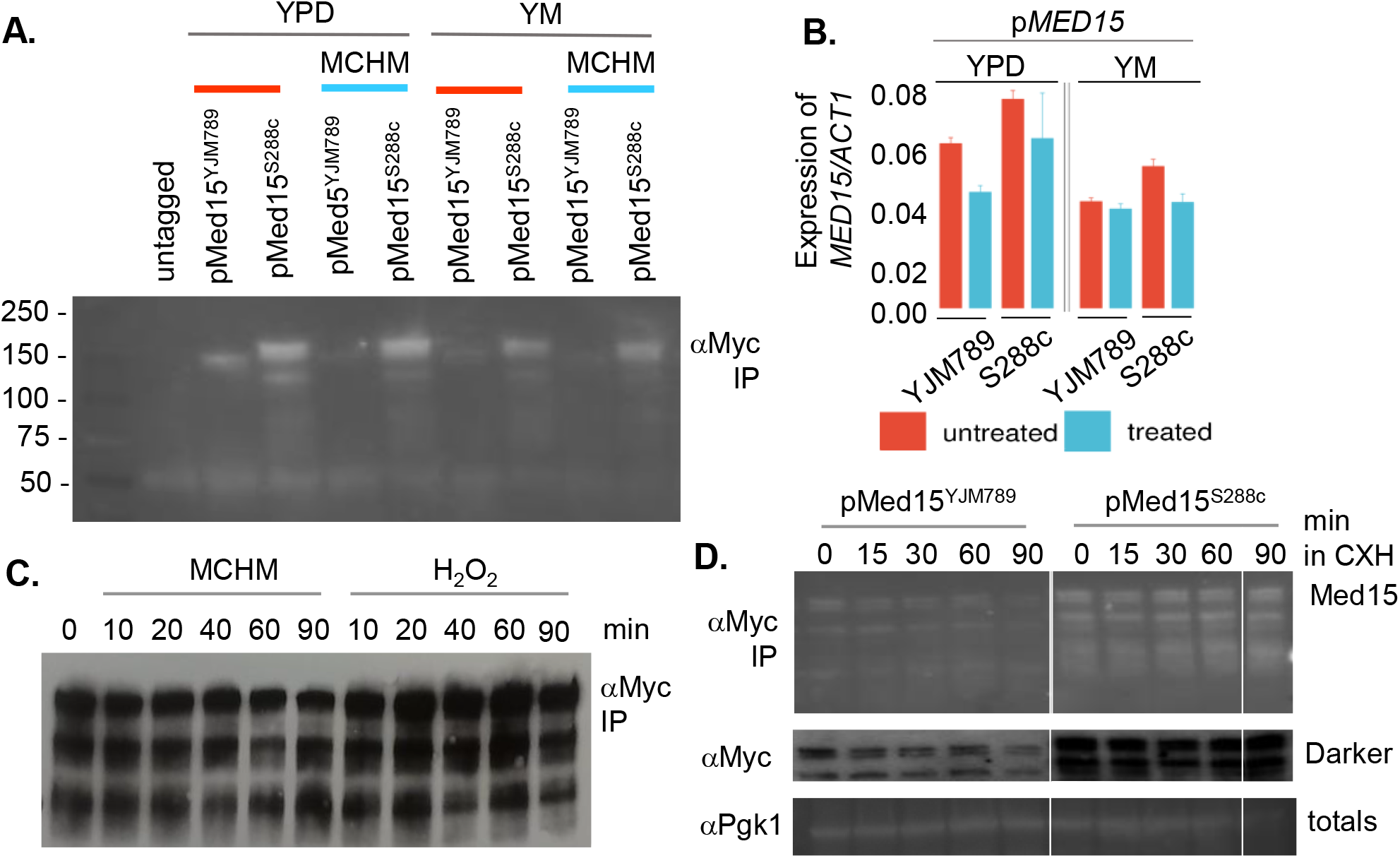
Changes in the gene expression levels of different alleles of Med15 treated with MCHM. **A.** Protein levels of Med15-13xMyc expressed from a plasmid in BY4741 *med15* yeast. Yeast were grown in selective media until mid-log and then shifted to 550 ppm MCHM for 90 minutes. Med15-13xMyc was immunoprecipitated from equal amounts of protein extract. **B.** mRNA levels of *MED15* expressed from a plasmid in BY4741 *med15* yeast normalized to *ACT1* mRNA. Transcript levels were extract from RNA-Seq data. Yeast were grown in YPD (with G418) or YM (yeast minimal media supplemented with HLM) and then treated with 550 ppm MCHM for 30 minutes. **C.** Western blot of Med15-Myc immunoprecipitated from BY4741 grown in YPD at 0, 10, 20, 40,60 and 90 minutes after the addition of 550 ppm MCHM or H_2_O_2_. **D.** Western blot of Med15-Myc immunoprecipitated from BY4741 carrying YJM789 and S288c alleles of Med15 from yeast grown in YPD at 0, 15, 30, and 90 minutes after the addition of cycloheximide. The total lysate was run separately and Pgk1 was blotted as a loading control.

Med15 contains multiple phosphorylations with the C-terminal MAD. It is unknown if these phosphorylations are regulated in a stress-dependent manner. Yeast expressing Med15^S288c^-Myc were treated with either MCHM or hydrogen peroxide for 90 minutes. There was no visible change in the pattern of Myc-tagged proteins in the western blot (Figure 3C). Next, the stability of the Med15 proteins was measured by treatment with cycloheximide, which blocks translation. While Med15^YJM789^ protein level was lower than Med15^S288c^ by the end of the time course, Med15^YJM789^ had decreased more relative to the levels of Med15^S288c^ (Figure 3D).

Med15 is important for response to many different stresses and to determine how genes were differentially regulated RNA-seq was carried out. BY4741 and its isogenic *med15* knockout were grown to log-phase and then treated with MCHM. In YPD, 149 genes were upregulated and 184 genes were downregulated in the *med15* knockout compared to BY4741 (Figure 4A and Table S2). The downregulated genes were related to metabolic processes of nucleosides and ribonucleosides, pyruvate metabolism, carbohydrates, and organophosphates catabolism, small molecule biosynthesis, oxidoreduction coenzyme metabolism, among others (Figure S2). In YPD, 46 GO terms were upregulated and 76 were down-regulated, while in YM, 35 were upregulated and 72 were downregulated. This set was not enriched in genes related to heat-shock response, drug/toxin transport, stress response, and cellular import as in (34) or in ribosome biogenesis as in (35). Sporulation related genes were upregulated (Figure S3), as previously reported in (34, 36); although, they are not yet known to have functional relevance in haploid cells. There were also genes involved in cell development, reproduction, morphogenesis and sulfur compound biosynthetic process. Previously study found that genes were upregulated sulfur metabolism in the *med15* mutant (34) also genes. When MCHM was added the differentially expressed genes increased in the *med15* mutants. 468 genes were upregulated and 278 were down regulated (Figure 4B and Table S2). Even with 90 more genes, there was extensive overlap in the functionality of the downregulated genes in the *med15* knockout compared to BY4741 in YPD only and YPD + MCHM, with only three more GO terms appearing: monosaccharide metabolism and organic acid and carboxylic acid biosynthesis (Figure S2). The difference was significant in the upregulated genes, not only in the number but in their functionality, as the GO terms overlap was low and a wide set of new terms related to ribosomes, polyamine transport and RNA export from nucleus appears. It is of note that in our study *med15* deletion caused the upregulation of ribosome biogenesis genes, contrary to the downregulation observed in (35). Also, their observed downregulation of this set of genes was the same in wild-type vs *med15* under osmotic stress, while we only observe the upregulation in the presence of MCHM, suggesting a fundamentally different mechanism of responding to osmotic stress and MCHM induced stress in yeast.

**Figure 4.**
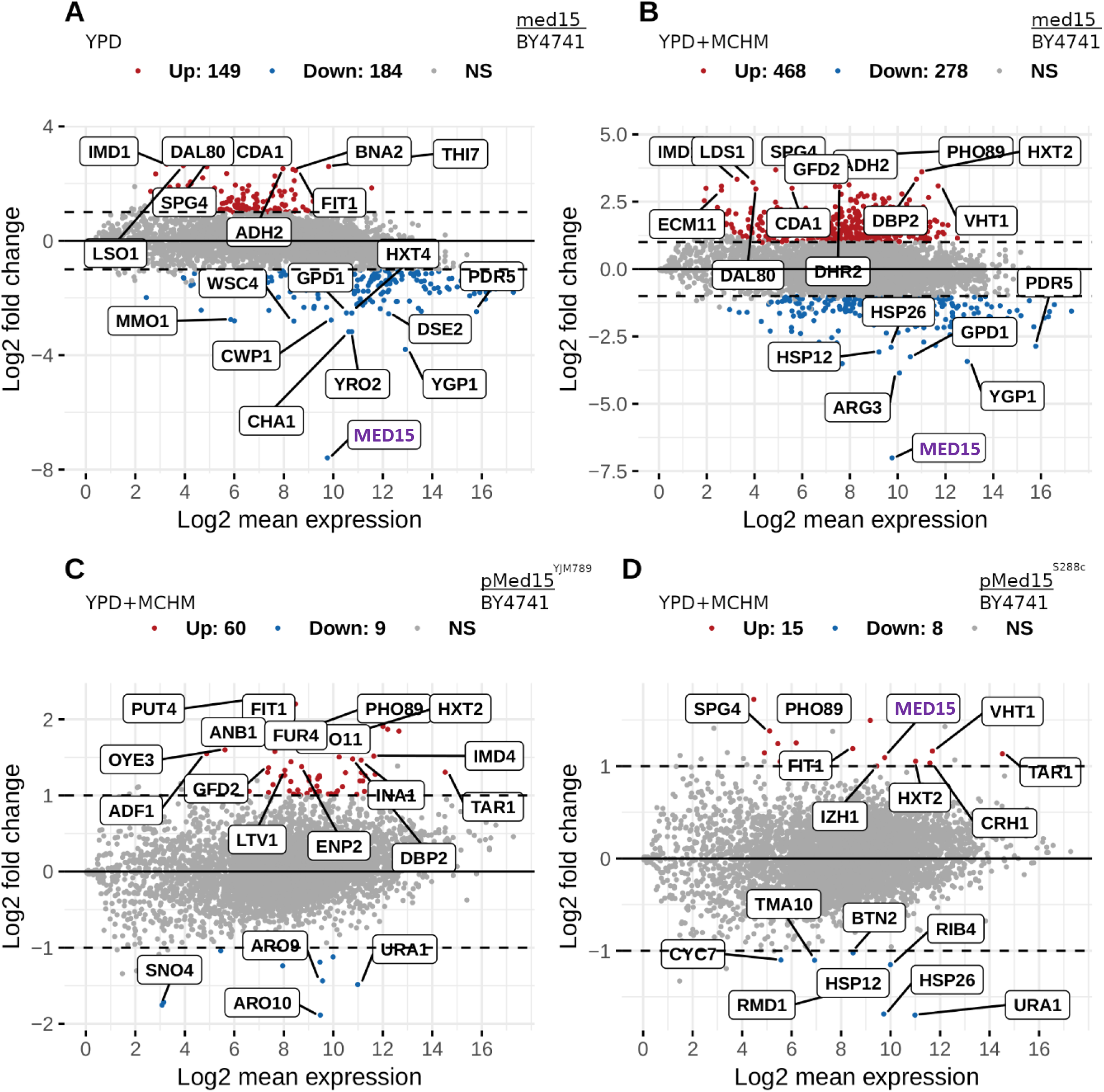
Changes in the transcriptome of BY4741 yeast carrying different alleles of Med15 treated with MCHM grown in YPD. **A.** Differentially expressed mRNA from wild-type yeast (BY4741) compared to a *med15* (*GAL11*) knockout strain grown in YPD. **B.** Differentially expressed mRNA from wild-type yeast (BY4741) compared to a *med15* (*GAL11*) knockout strain grown in YPD then shifted to 400 ppm MCHM for 30 minutes displayed on a log scale. **C.** Differentially expressed mRNA from wild-type yeast (BY4741) compared to a *med15* (*GAL11*) knockout strain carrying Med15^YJM789^ expressed from a plasmid grown in YPD then shifted to 550 ppm MCHM for 30 minutes. **D.** Differentially expressed mRNA from wild-type yeast (BY4741) compared to a *med15* (*GAL11*) knockout strain carrying Med15^S288c^ expressed from a plasmid grown in YPD with G418 then shifted to 550 ppm MCHM for 30 minutes.

By directly comparing the Med15^YJM789^ and Med15^S288c^ effect on gene expression, Med15^YJM789^ changed the expression of 69 genes and Med15^S288c^ 23 genes compared to BY4741 when treated with MCHM in YPD (Figure 4 C and D, respectively, Table S2). Med15 was among the overexpressed genes in Med15^S288c^ vs BY4741 (log2FC ~1.1) and probably the change of expression of the other 22 genes was due to this. The functional impact may be minimal as no term came out the GO analysis (Figure S2 and S3). Eight out of the nine downregulated genes in Med15^YJM789^ vs BY4741 were involved in small molecule biosynthetic process (Figure S2). Besides ribosome biogenesis, there were upregulated genes related to rRNA processing, ribonucleoside and glycosyl compound biosynthetic processes and ion transport (Figure S3).

The change of the media (YM instead of YPD) provoked a significant change in gene expression variation among the different cases being compared (Figure S4 and Table S2). But, the functional analysis of downregulated genes was strikingly similar to the one yeast grown in YPD (Figure S3 and S6). The functional analysis of upregulated genes in YM showed a different picture, with *med15* knockout verses BY4741. GO terms were almost the same regardless the presence of MCHM and with three new GO terms appearing in *MED15*^*YJM789*^ vs BY4741: sulfate assimilation, cysteine biosynthesis, and secondary metabolism.

Med15 binds upstream of many genes (Dunn, Gallagher, and Snyder, unpublished). Three genes were chosen for further characterization, *PTR2, PUT4*, and *YDJ1*. Except in MCHM treatment in YM, the levels of *PTR2* were significantly decreased and the levels of *PUT4* significantly increased in Med15^YJM789^ with respect to Med15^S288c^ in all other conditions, while the levels of *YDJ1* expression remained the same (Figure 5A). The knockouts of these genes were conferred MCHM sensitivity in YPD. However, in YM, only the *ydj1* yeast strain was also sensitive to MCHM (Figure 5B). Swapping the Med15 alleles in the *ydj1* knockout had no effect on growth. The *ydj1* knockouts were slow growing in BY4741 (Figure 5B and C) and are lethal in W303 (Caplan and Douglas 1991). The impact of the loss of Ydj1 on Med15 protein levels was measured by western blot (Figure 5D). The Myc tagged proteins isoforms were more heterogeneous in size in the *ydj1* mutant and the levels of Med15^YJM789^ increased to match that of Med15^S288c^. The slowest migrating band of Med15^YJM789^ increased to match that of Med15^S288c^.

**Figure 5.**
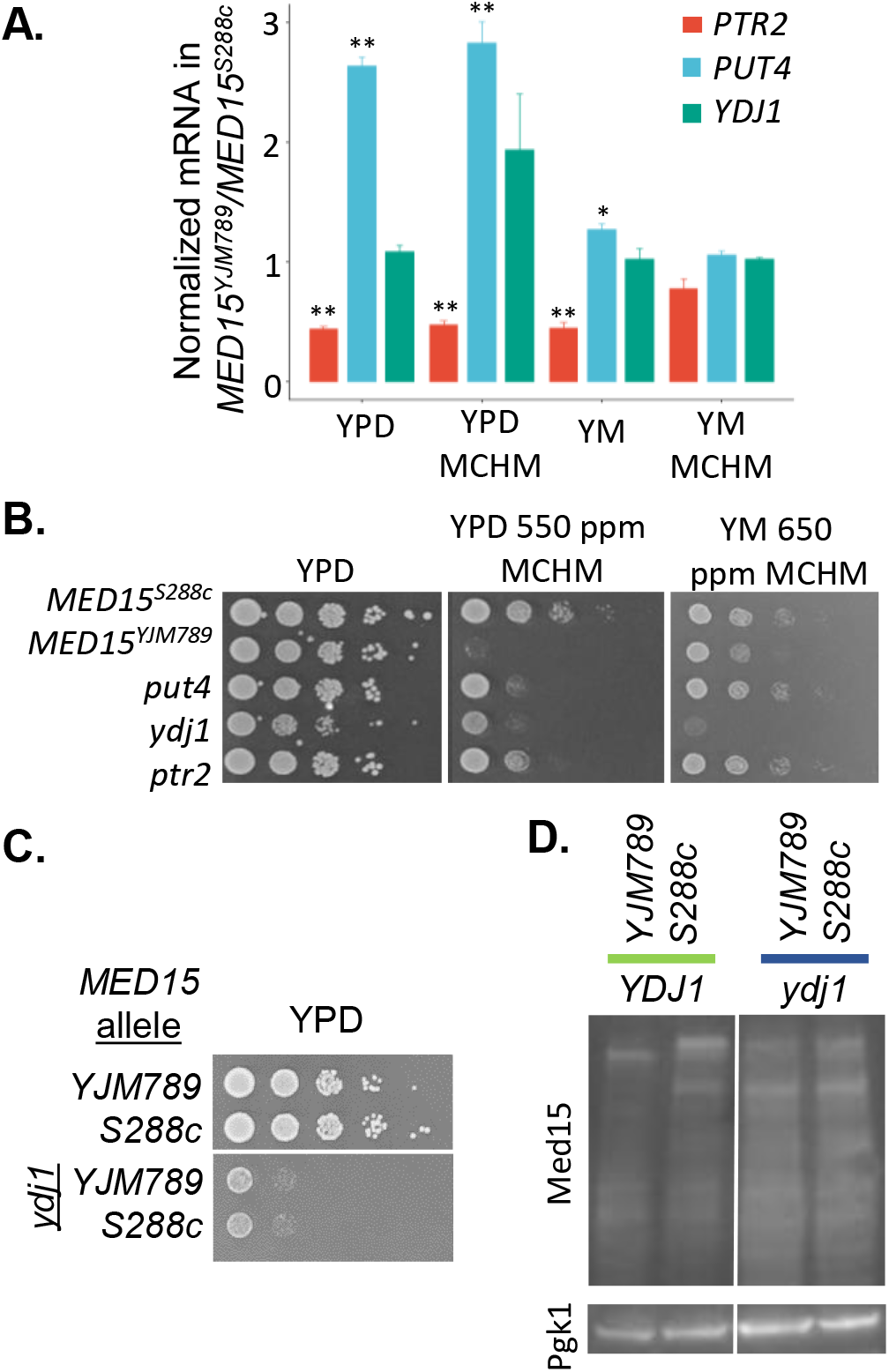
Conditions that affect the stability of Med15. **A.** Expression levels of *PTR2, PUT4*, and *YDJ1* extracted from RNA-seq data from supplemental **B.** Plasmids containing Med15^YJM789^ and Med15^S288c^ were transformed into double mutants of *med15* and *ydj1* in the BY4741 background. Serial dilutions of yeast on YPD were grown for 2 days at 30°C and then photographed. **C.** Serial dilution of yeast knockouts of *ydj1* yeast expressing YJM789 or S288c alleles of Med15-Myc on YPD. **D.** Western blot of YJM789 or S288c alleles of Med15-Myc immunoprecipitated from BY4741 or the *ydj1* mutant which were grown in YPD.

Med15 is nestled between Med16 and Med5 at the very distal end of the tail and Med2 and Med3, linking the tail to the C-terminal end of Med14 of the Mediator body (Figure 1B). Med15 alleles were swapped in *med2, med3*, and *med5* mutants and growth tested (Figure 6A). Overall the *med2* and *med3* mutants were slow growers in all conditions tested. However, the BY4741 *MED15*^*S288c*^ *med5* mutant grew similar to BY4741 *MED15*^*S288c*^ While the BY4741 *med5* mutants with either *MED15* allele grew the same. The loss of Med5 presumably does not alter Med15 binding to the rest of the Mediator and rescued the MCHM induced growth inhibition conferred by the Med15^YJM789^ allele while the loss of Med2 or Med3 causes similar growth defects. Like the *ydj1* mutants, both *med2* and *med3* mutants failed to grow in YM with MCHM. The levels of Med15 alleles were measured by western blot of immunoprecipitated Myc tagged proteins in YPD only because the mutants already had a growth phenotype (Figure 6B). The isoforms in the *med2* and *med3* mutants were similar to each other for each allele of Med15. The isoforms were numbered. The slowest migrating isoform 1 of Med15^S288c^ decreased in intensity while it increased in Med15^YJM789^. The second isoform that dominants wild type yeast with the Med15^YJM789^ remains in the *med2* and *med3* mutants. Med2 and Med3 are proposed to form a heterodimer independent of the Mediator (Béve *et al.* 2005). In cryo-EM structure, MAD which comprises the C-terminal tail of Med15 makes extensive contacts with Med2, Med3, Med4, Med5, and Med16 (2, 6). Med15 failed to immunoprecipitate other components of the Mediator in *med2* mutants and *vice versa* (Myers *et al.* 1999; Zhang *et al.* 2004).

**Figure 6.**
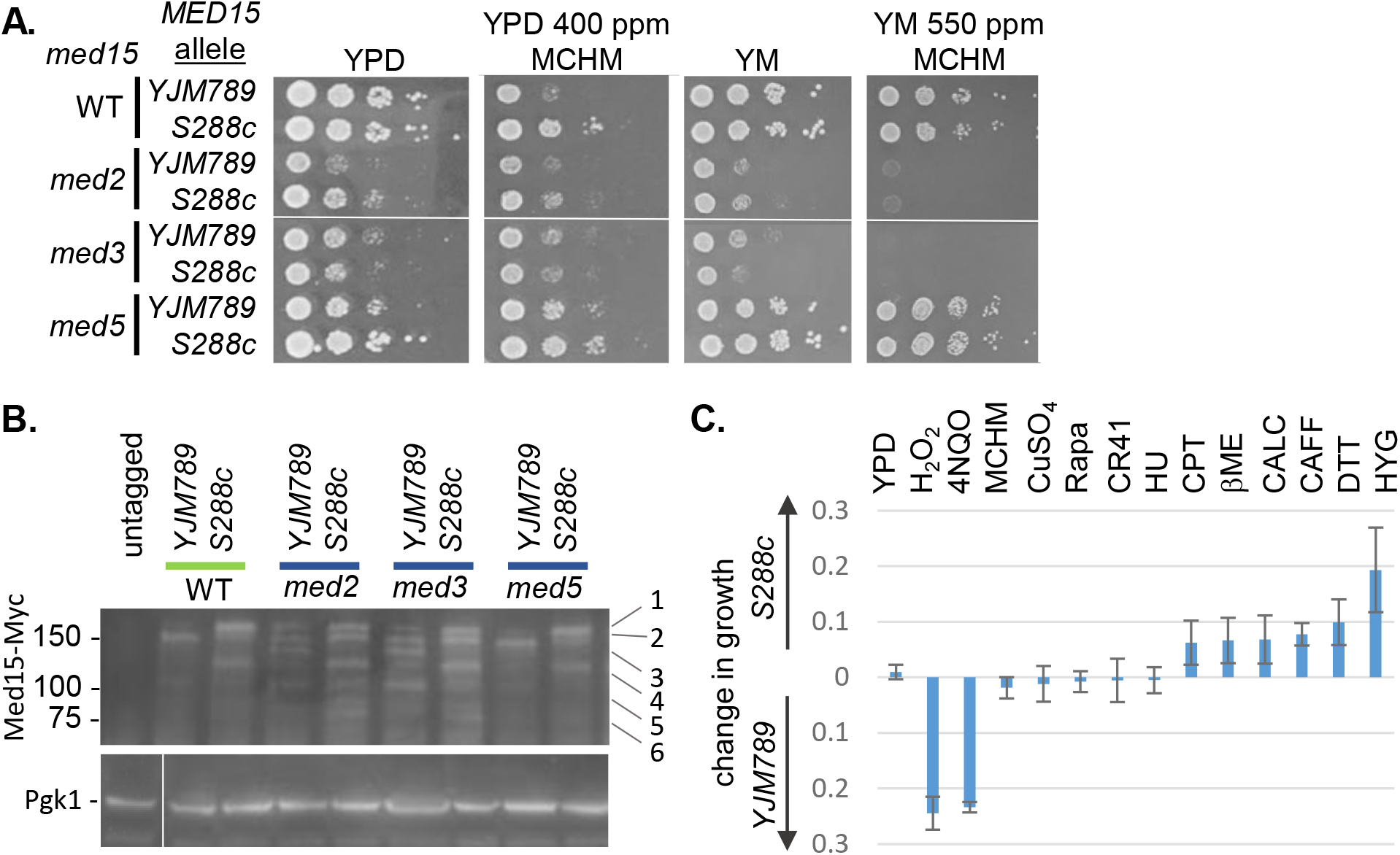
Impact of Mediator tail on different Med15 alleles. **A.** Serial dilutions of BY4741 with different alleles of Med15 combined with different mutants of the Mediator tail. **B.** Western blot of YJM789 or S288c alleles of Med15-Myc immunoprecipitated from BY4741 grown in YPD with *med2, med3* or *med15* deleted. Total lysate was run separately and Pgk1 was blotted as a loading control. **C.** Quantitative growth assays of BY4741 *med15* carrying different alleles of Med15. At the time where there was maximum growth difference, the OD600 of yeast carrying Med15^S288c^ was subtracted from OD_600_ of yeast carrying Med15^YJM789^. The following chemicals were added hydrogen peroxide (H_2_0_2_), 4-Nitroquinoline 1-oxide (4NQO), 4-Methylcyclohexanol (MCHM), copper sulfate (CuSO_4_), rapamycin (Rapa), glyphosate (CR41), hydroxyurea (HU), beta-mercaptoethanol (βME), calcofluor white (CALC), caffeine (CAFF), dithiothreitol (DTT), and hygromycin (HYG).

The two alleles of Med15 conferred different phenotypes not only against MHCM but also other chemicals (Figure 6C). Yeast with Med15^YJM789^ had greater resistance against compounds that generate free radicals directly, such as hydrogen peroxide and 4NQO which generates free radicals as it is metabolized (Rong-Mullins *et al.* 2018). MCHM is a volatile compound and when quantitative growth assays were carried out in small volumes, the MCHM evaporates (Gallagher *et al.* 2015) before the end of the growth assay. The Med15^S288c^ allele conferred resistance to reducing agents that cause unfold protein response such as beta-mercaptoethanol and DTT, DNA damaging chemicals such as camptothecin, and hygromycin which inhibits translation. Yeast with Med15^S288c^ were also more resistant to caffeine which in part can mimic the effects of TOR1 inactivation but not to rapamycin which also inhibits TOR1. Other chemicals that did not differentially inhibit yeast with different Med15 alleles were Credit41 (a commercial formation with glyphosate) and hydroxyurea which arrests cells in S phase by depleting nucleotides.

## Discussion

PolyQ expansion proteins were discovered to be the cause of numerous neurodegenerative diseases. Slippage of the DNA polymerase during DNA replication causes expansion and contraction of the repeats. In Huntington’s disease, expansions over 30 repeats are considered pathogenic and induce aggregation of Huntington protein. The two polyQ tracts in Med15 vary between 12 and 27 repeats. Changes in polyQ tracts of Med15 changed the response to numerous chemicals. Throughout the tail proteins of the Meditator, there is genetic variation that has yet to be explored. Reciprocal hemizygosity of *med15* mutants did not differentiate between the YJM789 or BY4741 alleles. Both hemizygous mutants were more sensitive to MCHM than the homozygous mutant, despite the YJM789 strain having a higher tolerance to MCHM than BY4741. Allele swapping of Med15^YJM789^ into BY4741 background conferred MCHM sensitivity. While in YJM789, expression of Med15^S288c^ did not change MCHM resistance. In both BY4741 and YJM7879 strains, the *med15* mutants were slow growing which was not affected at higher concentrations of MCHM. Making it appear that at the highest concentrations of the MCHM, the *med15* mutants were resistant to MCHM. MCHM sensitivity induced by expression of Med15^YJM789^ in BY4741 was not dominant. Therefore, we concluded that the YJM789 Mediator complex can better tolerate Med15 with a shorter polyQ tracts. While the BY4741 Mediator is incompatible with Med15 containing the expansion of the polyQ tracts. *MED15*^YJM789^ was expressed at slightly lower levels and the protein were even lower levels than Med15^S288c^ and was also less stable. While changes in mRNA levels contribute to lower protein levels, the longer polyQ tracts in Med15^YJM789^ may also slow translation or increased ubiquitin-dependent degradation as the protein is less stable when translation was inhibited.

Ydj1 was required for stability of Med15 protein and it was difficult to assess the role of Ydj1 on Med15 protein stability because of the extremely slow growth of the *ydj1* mutants. Ydj1 also has a role at H3 histone eviction when transcription is induced. In one example, Gcn4 binding to promoters was not reduced in a *yjd1* mutant (Qiu *et al.* 2016) or at the *GAL1* promoter (Summers *et al.* 2009). Hsp70 associates with several Hsp40-like proteins including Ydj1, a type 1 Hsp40, that stimulates Hsp70 activity. Ydj1 is localized to the perinuclear and nuclear membranes (Caplan and Douglas 1991). The role in nucleosome eviction may be indirect by helping to fold Med15. Ydj1 can inhibit the SDS-resistant aggregation of the polyQ containing a fragment of Htt in yeast (Krobitsch and Lindquist 1999; Muchowski *et al.* 2000). The Med15 fragment containing the polyQ aggregates *in vivo* (Zhu *et al.* 2015) as well as when full-length Med15 is overexpressed. Ydj1 was required for both alleles of Med15 protein stability as the isomers that were Myc staged became less distinct maintaining Med15^YJM789^ true to size compared to Med15^S288c^.

Med2, Med3, and Med15 can be recruited to chromatin independent of the rest of the Mediator complex (45). From the recent structures of the Mediator, Med2 and Med3 bind the C-terminal tail of Med14 in the body and directly binds Med15. Med15, in turn, binds Med16 then Med5 is at the very distal end of the tail. Double mutants of *med15* with either *med2* or *med3* are viable while *med15* combined with *med5* or *med16* are lethal (36). The Med15-Med5-Med16 complex is posited to have an essential function independent of the full Mediator complex (36). By their nature, the structure of intrinsically disordered regions (IDR) are difficult to determine and are important for changes in protein complex conformations (19, 23, 46–48). The fuzzy/ IDR domains of Med15 and the expansions of the polyQ tracts increased phenotypic diversity. While Rim101, a transcription factor with a polyQ track and affects allele-specific expression in one strain background but not others tested (Read *et al.* 2016). There are multiple phosphorylations in Med15 that regulates transcriptional response to stress (35). Expression of the incompatible Med15 allele changed the response to MCHM as other polymorphic transcription factors change the response to other chemical stressors (28). Variation in key regulators permits the expression of cryptic genetic variation to alter phenotypes. These proteins are master variators such as Yrr1 which has a single polymorphic phosphorylatable amino acid that switches the growth of yeast in the presences of 4NQO and nonfermentable carbon sources (28).

## Acknowledgments

Angela Lee generously gave us the yeast knockout collection. Michael Ayers and the Gallagher lab provided fruitful discussion. Noor Malik made strains and was funded through WVU Summer Undergraduate Research Experience. We would like to acknowledge the WVU Genomics Core Facility, Morgantown WV for the support provided to help make this publication possible including RNA-seq library construction and sequencing. The PGK1 antibody was a gift from Jeremy Thorner. This work was supported by a grant from the NIH (R15ES026811-01A1).

## Author contributions

JEGG designed the experiments and wrote the manuscript. AP carried out RNA-seq analysis. CN carried out yeast growth assays.

---

**Figure S1 A.** Reciprocal hemizygotes of Med15 in BY4741xYJM789 hybrids were grown on MCHM in YPD. Med15 was tagged at the chromosomal locus with 13xMyc at the C-terminal end or knockout with a dominant drug marker in haploid parents. Yeast were then mated, and diploids selected. An equal number of yeast were serially diluted and plated onto YPD with the indicated amount of MCHM **B.** BY4741 yeast (wildtype) and BY4741 *med15::NatR* (*med15Δ)* were transformed with pGS35 (empty) or pGS35-*MED15-Myc* (p*MED15*^*YJM789*^Myc and p*MED15*^*S2889*^Myc). Plasmids were maintained with G418 in YPD and YM with glutamate (MSG) as the nitrogen source. Yeast were serially diluted and plated with indicated amounts of MCHM.

**Figure S2** GO term analysis on genes that are downregulated in *med15* mutants grown in YPD or YPD + 550 ppm MCHM compared to BY4741 and BY4741 expressing *MED15*^*YJM789*^ compared to BY4741 grown in YPD + 550 ppm MCHM.

**Figure S3** GO term analysis on genes that are upregulated in *med15* mutants grown in YPDor YPD + 550 ppm MCHM compared to BY4741 and BY4741 expressing *MED15*^*YJM789*^ compared to BY4741 grown in YPD + 550 ppm MCHM.

**Figure S4.** Changes in the transcriptome of BY4741 yeast carrying different alleles of Med15 treated with MCHM grown in YM. **A.** Differentially expressed mRNA from wild-type yeast (BY4741) compared to a *med15* (*GAL11*) knockout strain grown in YPD. **B.** Differentially expressed mRNA from wild-type yeast (BY4741) compared to a *med15* (*GAL11*) knockout strain grown in YM then shifted to 550 ppm MCHM for 30 minutes. **C.** Differentially expressed mRNA from wild-type yeast (BY4741) compared to a *med15* (*GAL11*) knockout strain carrying Med15^YJM789^ expressed from a plasmid grown in YM then shifted to 650 ppmMCHM for 30 minutes. **D.** Differentially expressed mRNA from wild-type yeast (BY4741) compared to a *med15* (*GAL11*) knockout strain carrying Med15^S288c^ expressed from a plasmid grown in YM with G418 then shifted to 650 ppm MCHM for 30 minutes.

**Figure S5** GO term analysis on genes that are downregulated in *med15* mutants grown in YPD or YM + 650 ppm MCHM compared to BY4741 and BY4741 expressing *MED15*^*YJM789*^ compared to BY4741 grown in YPD + MCHM.

**Figure S6** GO term analysis on genes that are upregulated in *med15* mutants grown in YPD or YM + 650 ppm MCHM compared to BY4741 and BY4741 expressing *MED15*^*YJM789*^ compared to BY4741 grown in YM + MCHM.

**Table S1:**
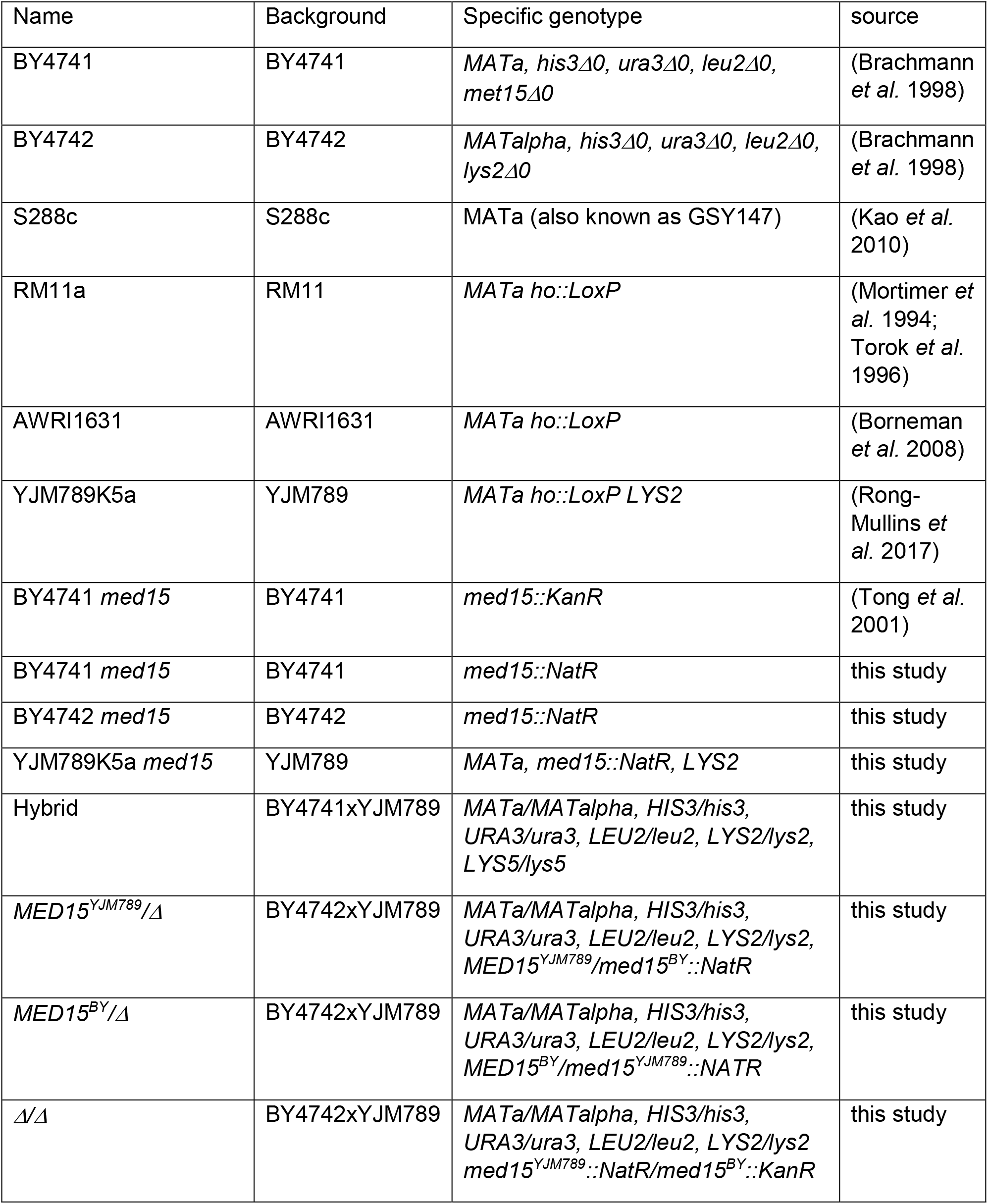
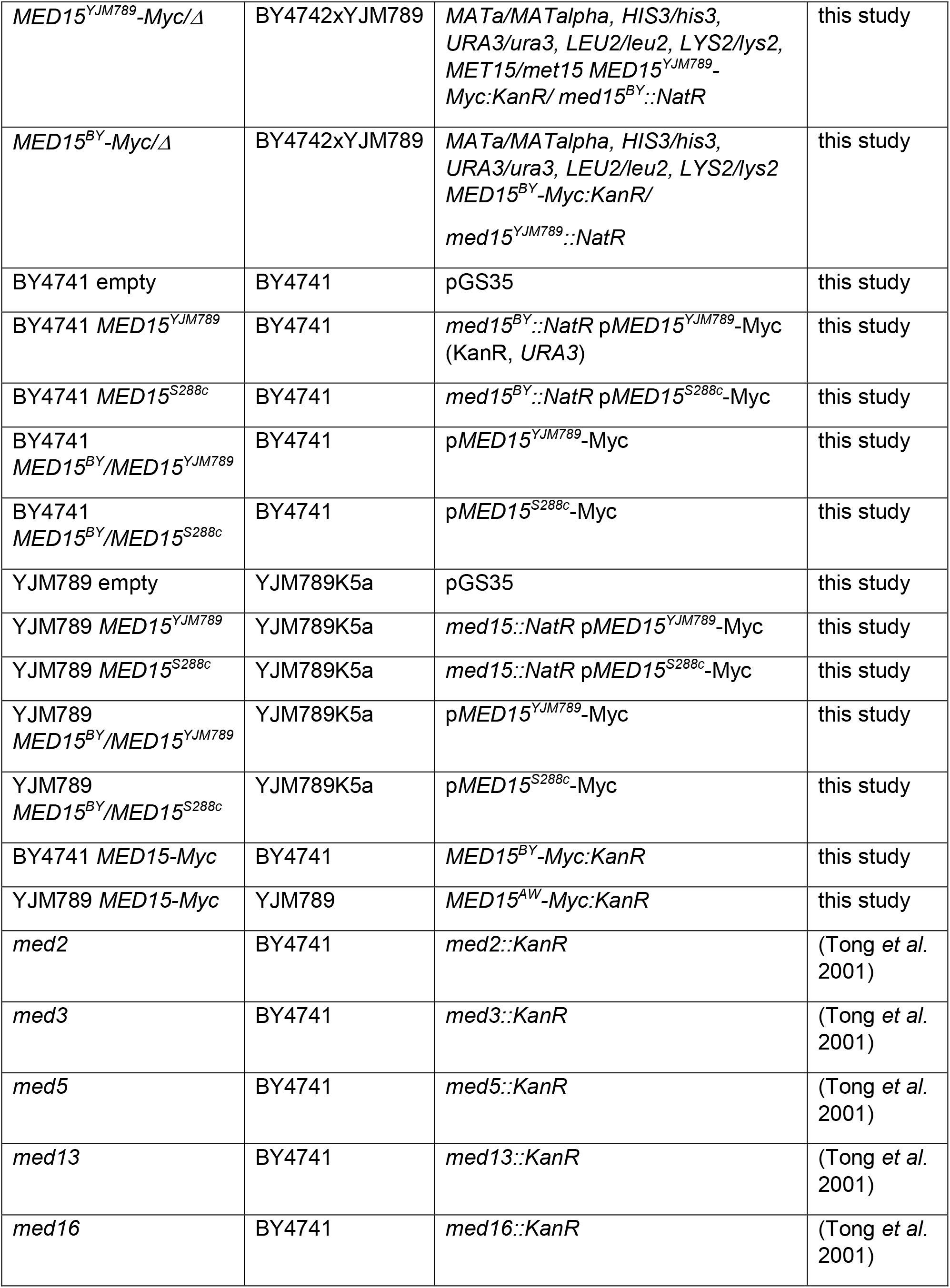
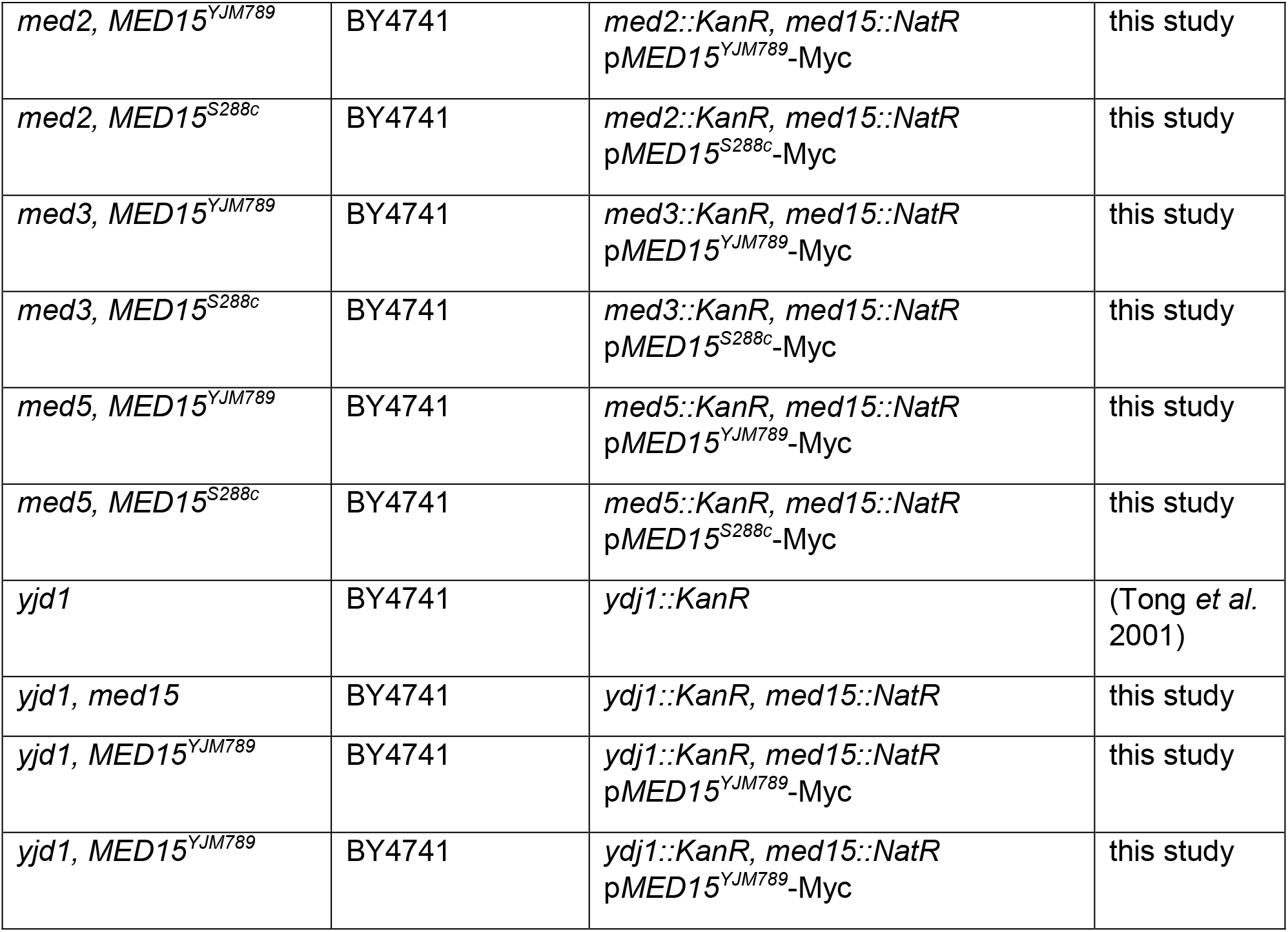
Strain list

**Table S2** Differentially Expressed Gene list from BY4741, BY4741 *med15::NAT* (BYmed15), BY4741 *med15::NAT* with pMed15^YJM789^-Myc (BYpMM_789), and BY4741 *med15::NAT* with pMed15^S288c^-Myc (BYpMM_S288c) grown in YPD or YM with or without 550 ppm MCHM.

## Materials and Methods

### Strain construction

Med15 sequences were extracted from the sequenced genomes of BY4741, BY4742, AWRI1631, RM11-1a and YJM789 (Song *et al.* 2015). Med15 was tagged at the C-terminus with 13x Myc tag with KanR as the selectable marker in BY4741 and YJM789 as previously described (28, 59). Primers to the genomic *MED15* amplified 499 nucleotides upstream from the start (TGGCGGCCGCTAACAAGCAATTACATATTCC) and a 3’ tagging primer to including the promoter, coding region, Myc tag and KanR marker (AGTAACTTCAAAAGTATCAAAAGTATGGAAACTTCAAATGTTTACGACTCACTATAGGGA). The PCR product was then cloned into the NotI restriction site in pRS316. *MED15* was knocked out in YJM789K5a (isogenic with YJM789 except as a MATa prototroph) and then backcrossed to generate YJM789K6alpha as previously described (Rong-Mullins *et al.* 2017). BY4741 knockout yeast of *med2, med3, med5*, and *ydj1* (Giaever *et al.* 2002) were crossed with BY4742 *med15::NatR* to generated double mutants and then transformed with plasmids containing different alleles of *MED15*.

### Growth Conditions

Plasmids were maintained with the addition of 0.5 mg/ml G418 in YPD. In minimal media (YM) plasmids were maintained by supplementing media with uracil, histidine, and methionine or by switching the nitrogen source to glutamate (MSG), and then adding G418 with amino acids as needed. Yeast were grown in liquid media as indicated to mid-log phase and then 550 ppm MCHM was added to YPD (650 ppm was added to YM) and cells were harvested after 30 minutes of exposure. Western blots were carried out as previously described (Gallagher *et al.* 2014). Solid media plates were cooled to 65°C before MCHM was added and gently mixed until solution. Plates were used within 24 hours to limit evaporation of MCHM. Yeast were serially diluted 10-fold and spotted on to solid media. Plates were photographed after 2-3 days of growth. For multiple drug screening in the TECAN, the automated plate reader, yeast were grown to stationary phase and then diluted to 0.1 OD with appropriate drugs and read at OD_600_ (Rong-Mullins *et al.* 2017). The following chemicals were added 3mM hydrogen peroxide (H_2_0_2_), 0.25 μg/ ml 4-Nitroquinoline 1-oxide (4NQO), 400 ppm 4-Methylcyclohexanol (MCHM), 1 mM copper sulfate (CuSO_4_), 7.5 ng/ml rapamycin (Rapa), 0.1% glyphosate (CR41), 100 mM hydroxyurea (HU), 20 μg/ml camptothecin (CPT), 8.5 mM beta mercaptoethanol (βME), 5 mM calcofluor white (CALC), 2.5 mM caffeine (CAFF), 20 mM Dithiothreitol (DTT), and 50 μg/ ml hygromycin. Cells were grown with readings taken every hour. During log-phase, the OD_600_ of yeast carrying *MED15*^*S288c*^ was subtracted from *MED15*^*YJM789*^ at the point of maximal growth difference.

### Transcriptomics

RNA-seq was carried out in biological triplicate from yeast grown in YM supplemented with histidine, leucine and methionine or YPD with G418. PolyA RNA was selected using Karpa Stranded RNA-seq library preparation kit according to manufacturer’s instructions (catalog number KK8401). Libraries were sequenced on Illumina PE50bp high output flowcell. Basecalls were performed with Illumina’s FASTQ Generation (v1.0.0) available in BaseSpace. Transcripts quantification was done with salmon (v0.9.1) vs the transcripts file BY4741_Toronto_2012_cds.fsa (available from https://downloads.yeastgenome.org/sequence/strains/BY4741/BY4741_Toronto_2012/). This data is available from GSE, accession number GSE129898 (https://www.ncbi.nlm.nih.gov/geo/query/acc.cgi?acc=GSE129898). Quantification tables were imported to R (3.4.4) and gene level analysis was created with the tximport (1.6.0) package. For the transcripts to gene translation the homemade R package TxDb.Scerevisiae.SGD.BY4741 was used. This package was built from the BY4741_Toronto_2012.gff file using GenomicFeatures (1.30.3). The gene differential expression analysis and the data quality assessment were done with DESeq2 (1.18.1). p values were adjusted to an FDR of 0.005. The MA-plots were done with ggpubr (0.1.6).

GO term analysis was carried out with clusterProfiler (61) (3.6.0). The ORF names from genes up or downregulated in each condition were translated to the correspondent Entrez id using the function bitr and the package org.Sc.sgd.db. The resulting gene clusters were processed with the compareCluster function, in mode enrichGO, using org.Sc.sgd.db as a database, with Biological Process ontology, cutoffs of p-value = 0.01 and q value = 0.05, adjusted by FDR, to generate the corresponding GO profiles, which were then simplified with the function simplify. The simplified profiles were represented as dotplots, showing up to 15 more relevant categories.

### Western blot

Proteins were extracted, immunoprecipitated, separated in 5-12% SDS-PAGE, and transferred onto 0.2 micron PVDF as previously described (28). Antibodies were diluted into freshly made 3% BSA Fraction V in TBS-Tween. ECL kit and HRP secondary antibodies were used to visualize mouse anti-Myc E910 (1:7,500) from various manufacturers and rabbit anti-PGK (1:10,000) on a Protein Simple using default chemiluminescence setting.

