## Supplemental Figures for "The polymorphic PolyQ tail protein of the Mediator Complex, Med15, regulates variable response to stress"

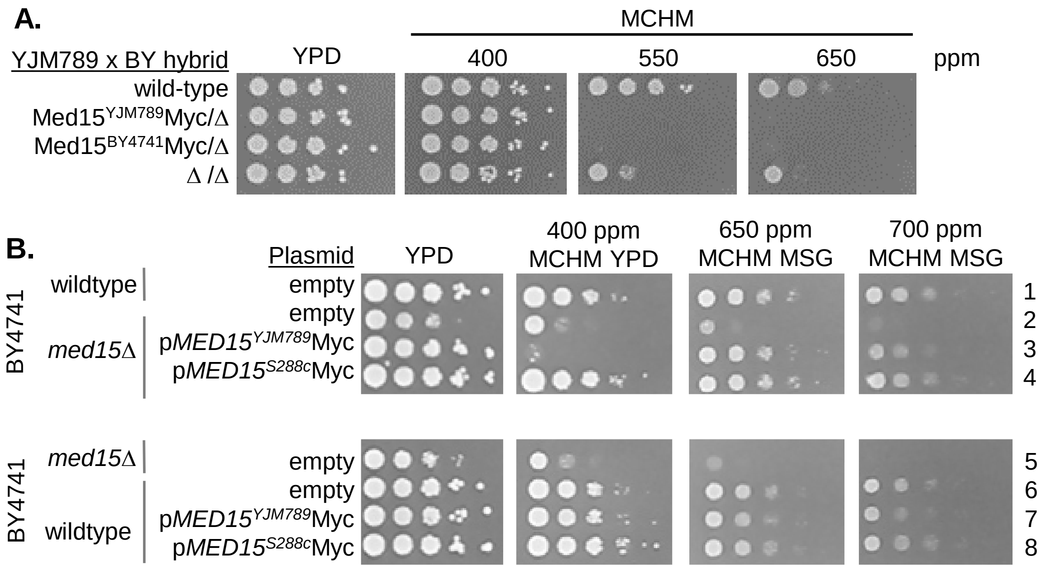


**Figure S1 Gallagher**

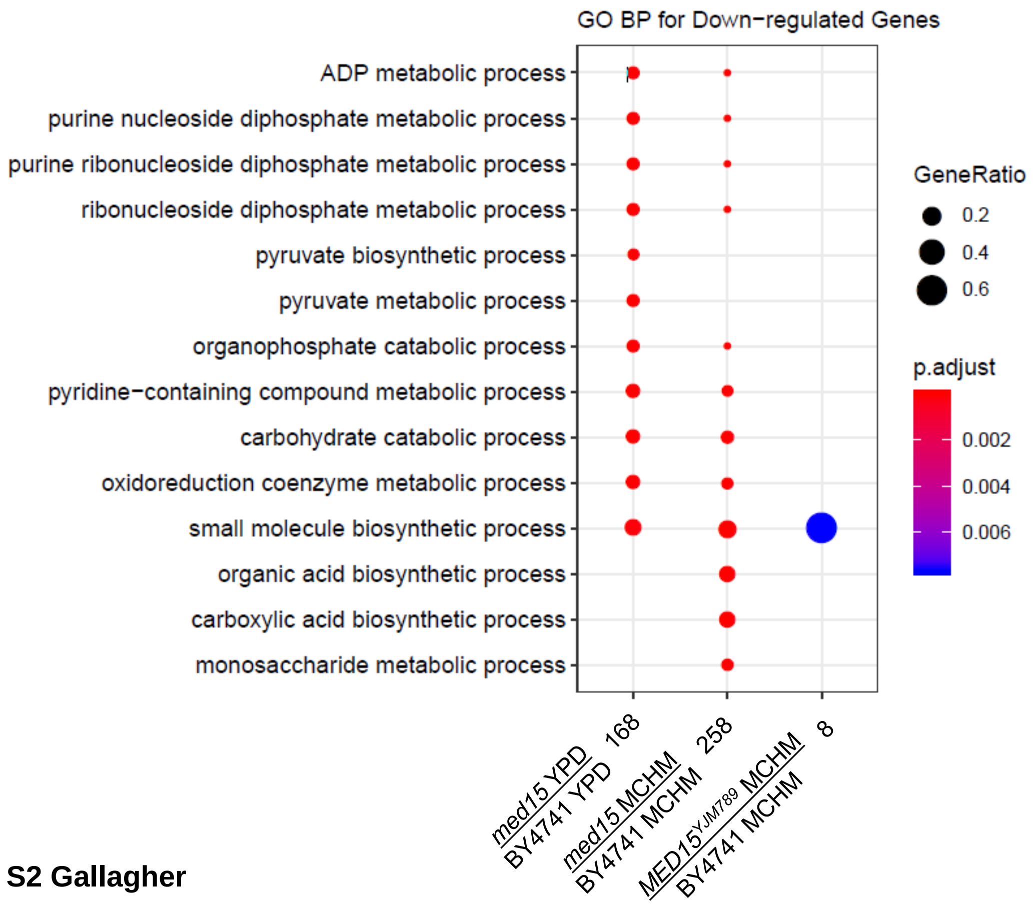


**Figure S2 Gallagher**

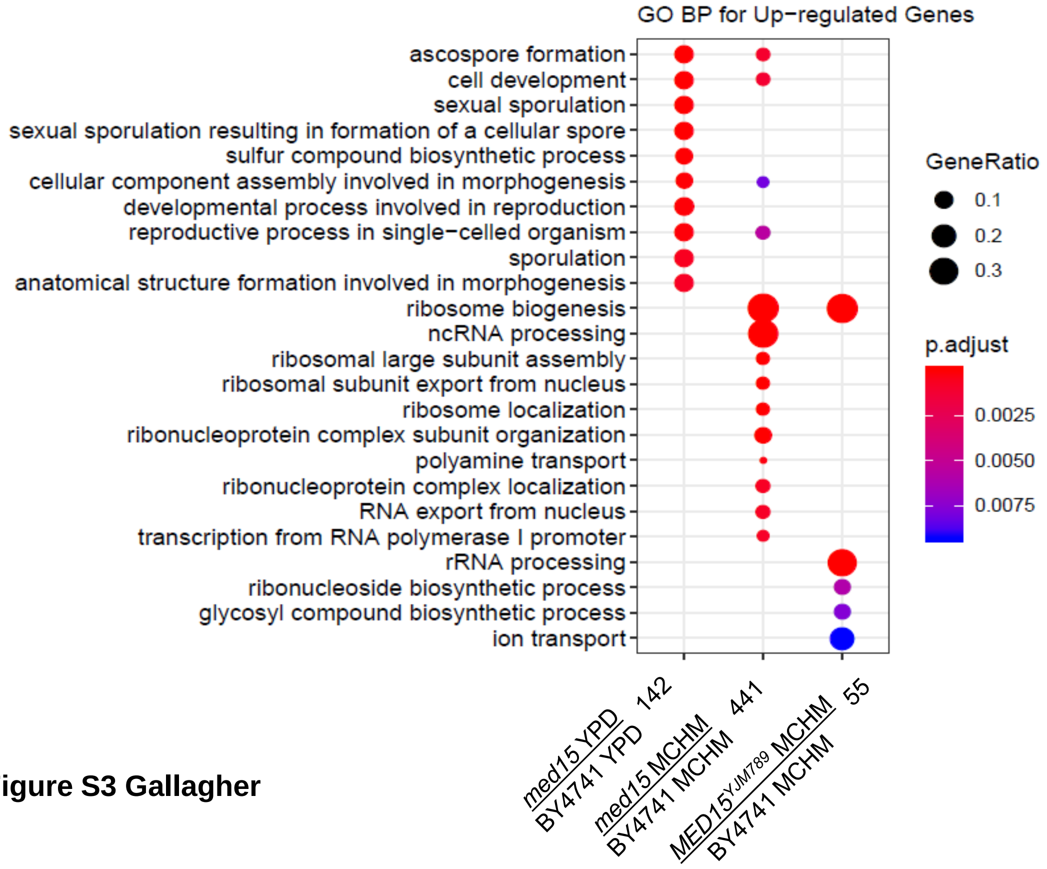


**Figure S3 Gallagher**

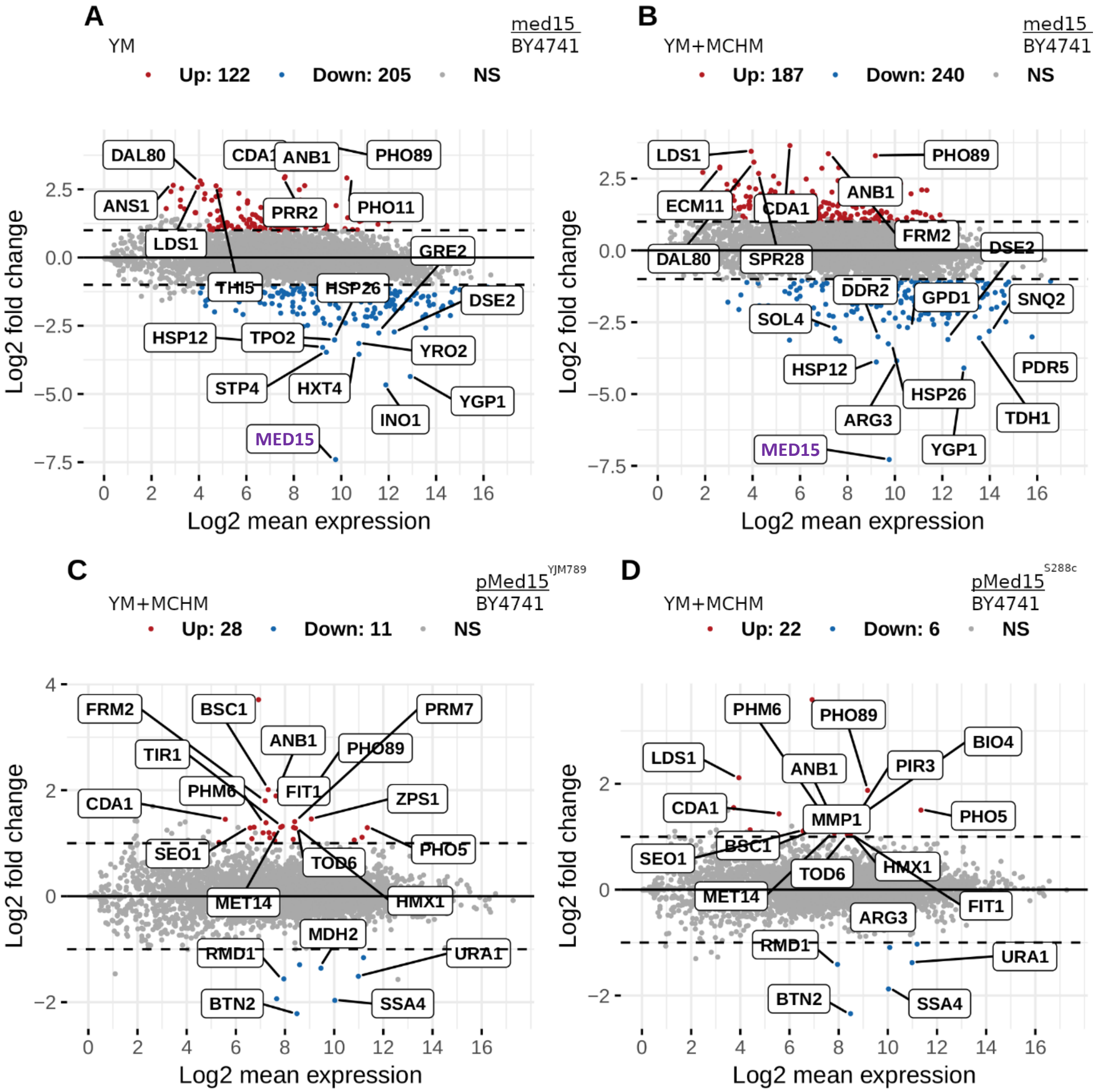


**Figure S4 Gallagher**

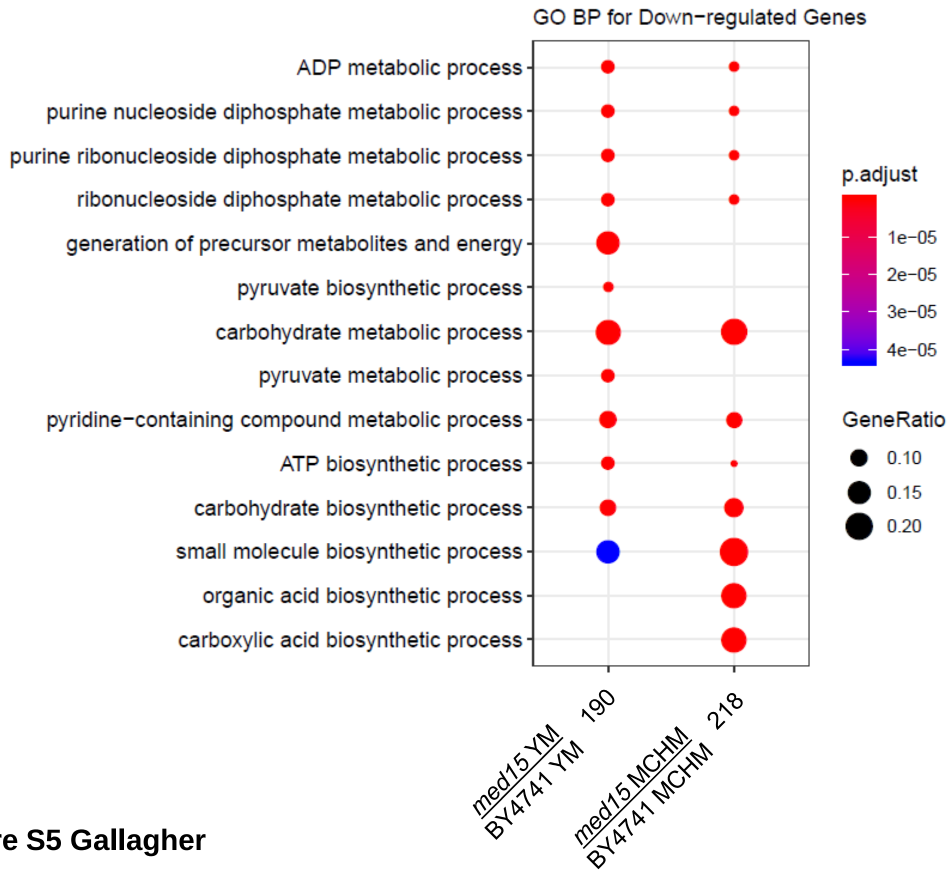


**Figure S5 Gallagher**

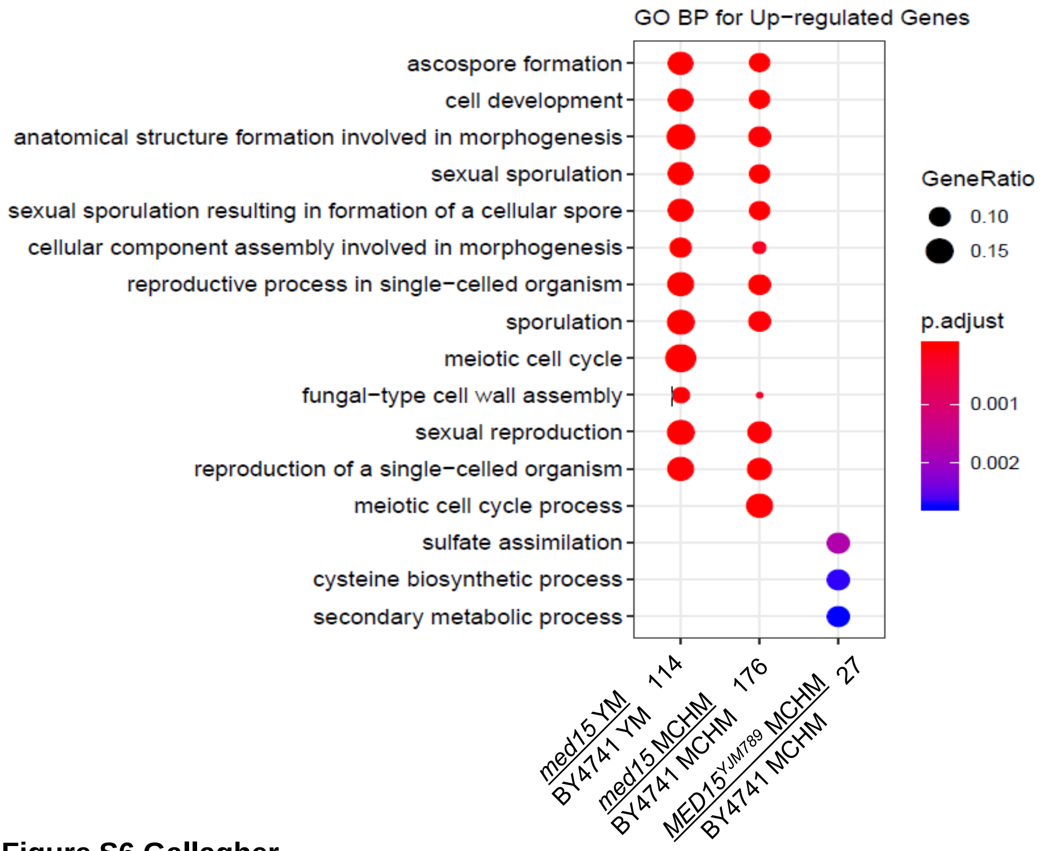


**Figure S6 Gallagher**
